## Supplementary Results for "Highly multiplexed ddPCR-amplicon sequencing reveals strong *Plasmodium falciparum* population structure and isolated populations amenable to local elimination efforts in Zanzibar"

**Supplementary Figures**

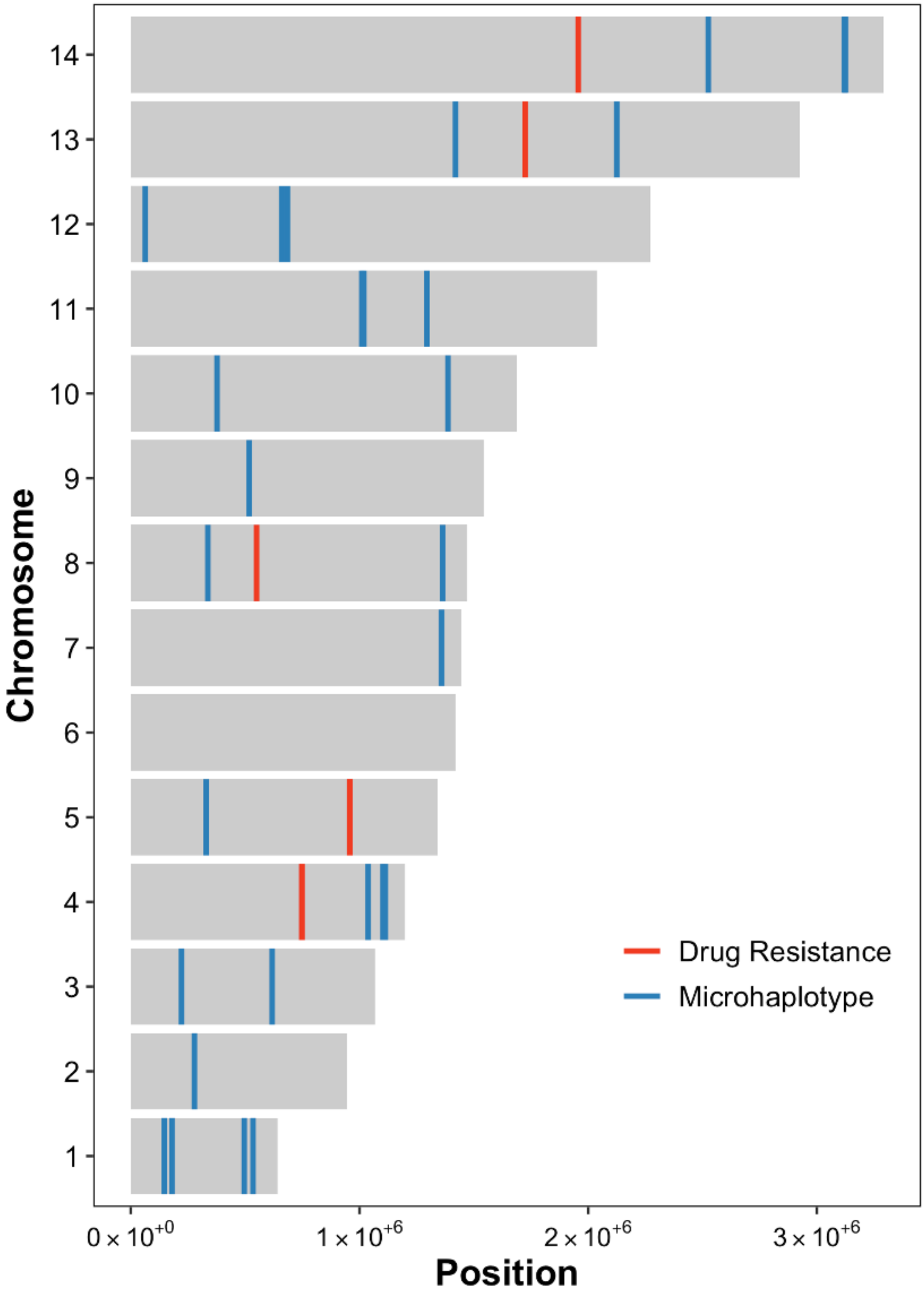


***Figure S1**. Chromosomal location of microhaplotypes (blue) and drug resistance loci (red) in the P. falciparum genome.*

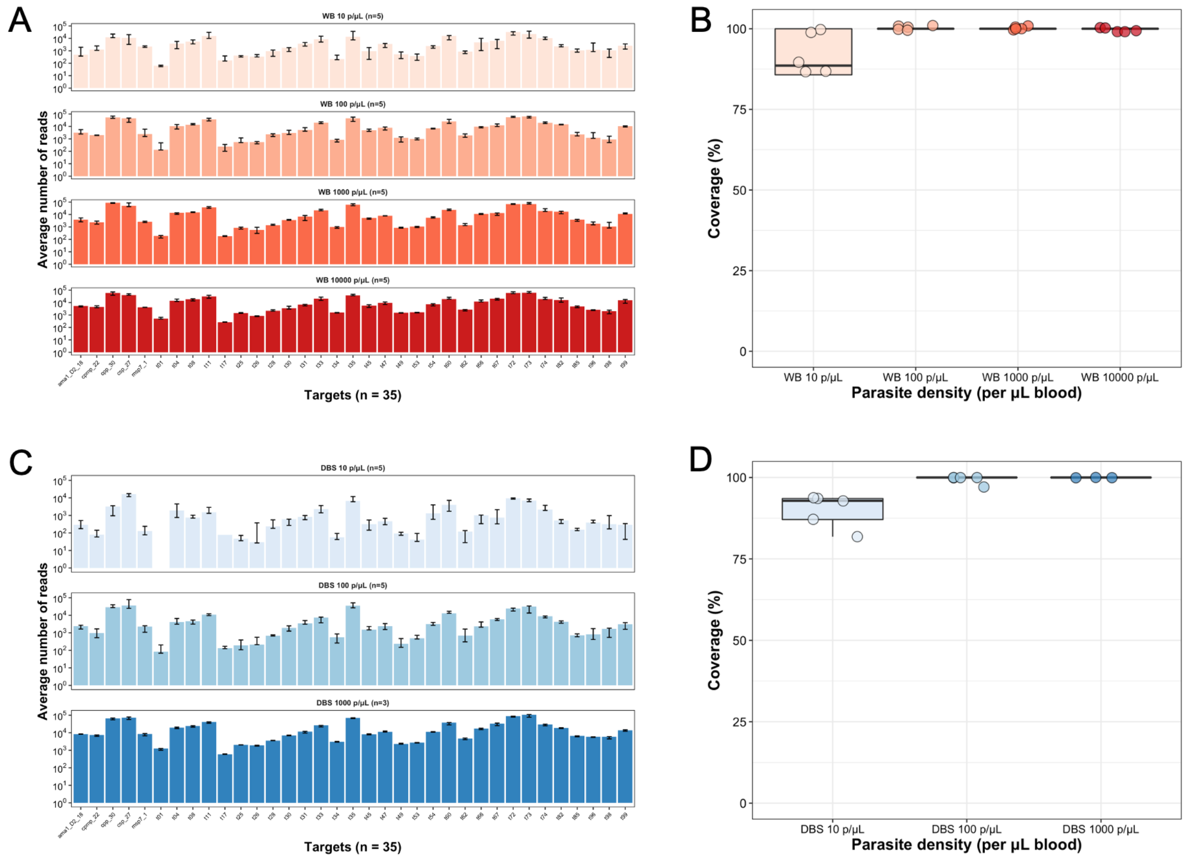


***Figure S2***. *Evenness and coverage of multiplexed amplicon sequencing of microhaplotypes (n=28) and drug resistance loci (n=7).* ***A & C***, *Average number of reads per target per sample. The median (bars) and interquartile range (error bars) are shown.* ***B & D***, *Boxplot summarizing the coverage of microhaplotype loci and drug resistance targets by parasite density. Samples are 3D7 whole blood (reds) ranging from 10,000 – 10 parasites/µL or 3D7 DBS (blues) ranging from 1,000 – 10 parasites/µL.*

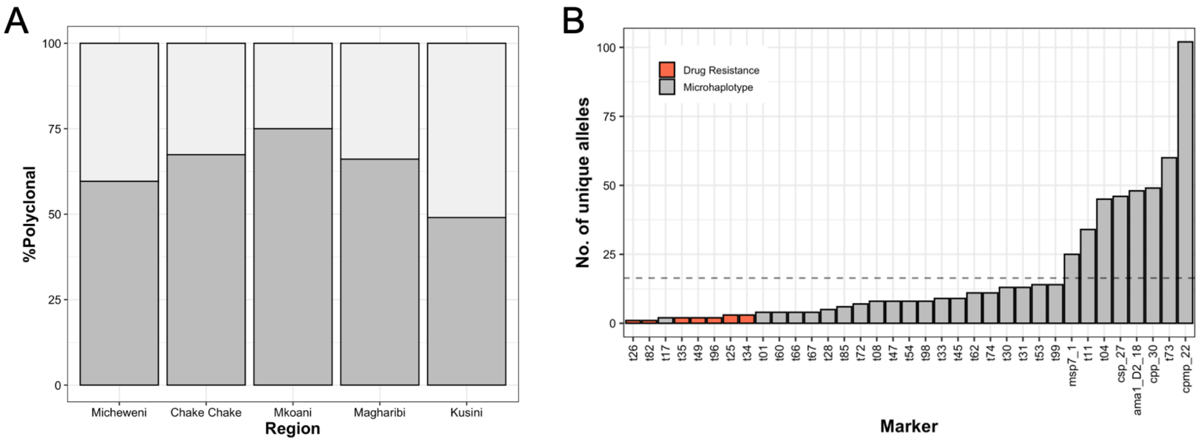


***Figure S3. A,*** *Percentage polyclonal infections by the 5 districts in Zanzibar (chi-square statistic: 6.49; p = 0.166).* ***B,*** *Number of unique alleles of microhaplotypes and drug resistance loci in all 290 Zanzibar samples (mean A: 16.7, range: 1-102). Dashed line indicates mean A.*

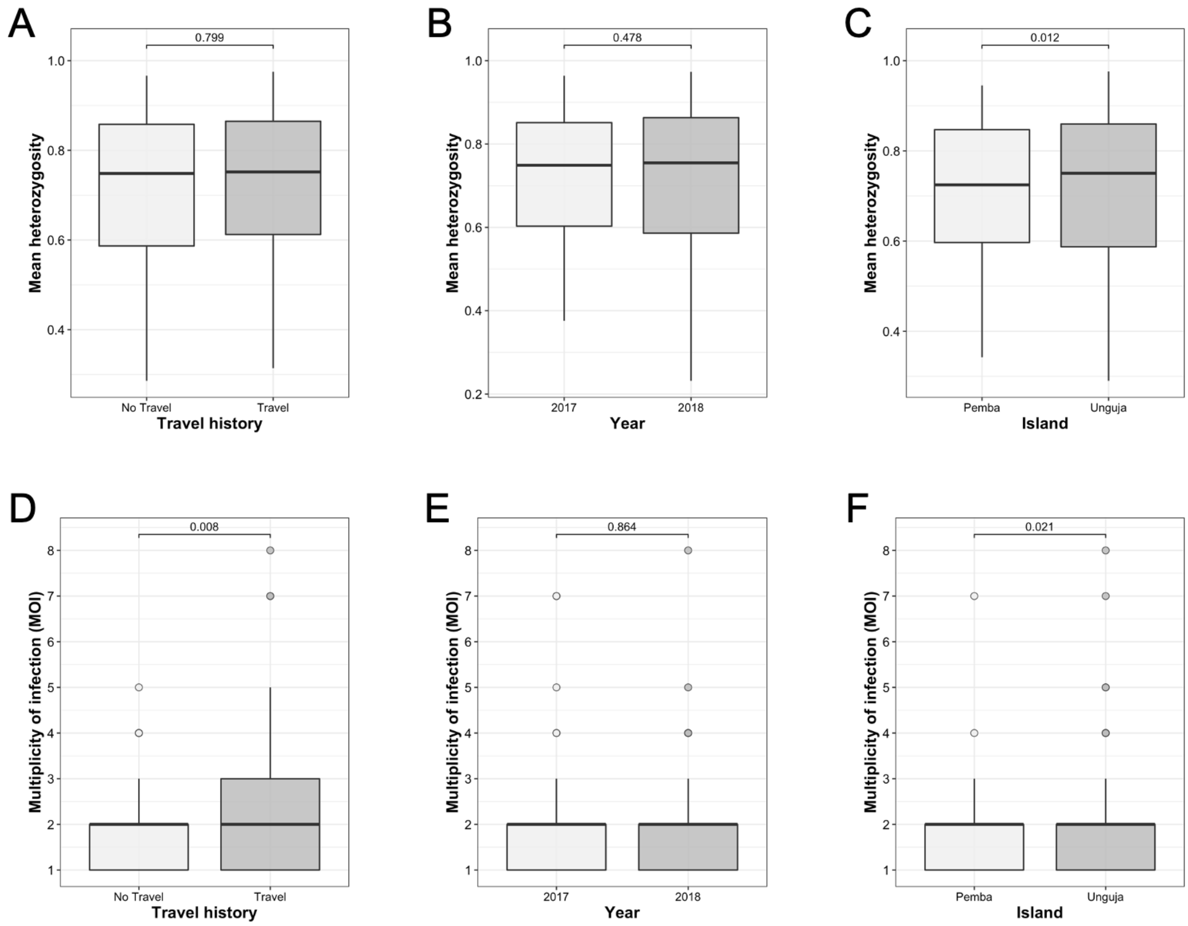


***Figure S4.*** *Multiplicity of infection and population diversity by travel history, year, and island.* ***A, B & C****, Population diversity measured as the distribution of heterozygosity in 28 microhaplotypes. Bonferroni adjusted pairwise p-values are indicated.* ***D, E & F****, Multiplicity of infection (MOI). Bonferroni adjusted p-values are indicated.*

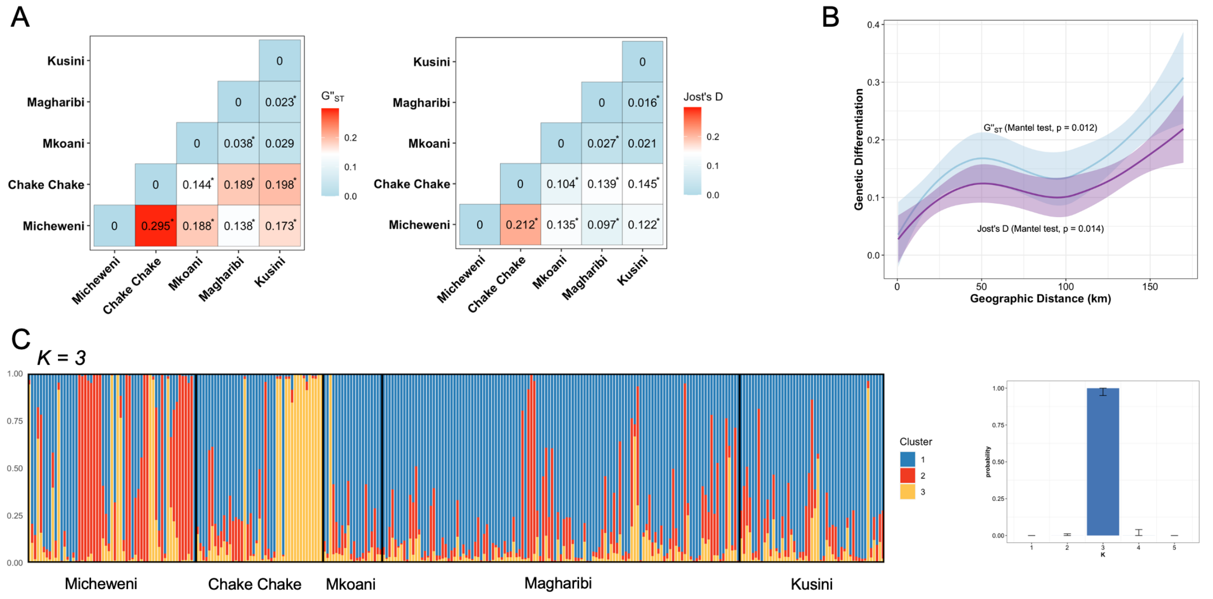


***Figure S5***. *Genetic differentiation and population structure.* ***A,*** *Pairwise* $G''_{ST}$ *(left) and Jost's D (right) between districts (* indicates P < 0.01; permutation test (100,000 permutations))* ***B****, Relationship between pairwise genetic differentiation and geographic distance in Zanzibar between 29 shehia pairs (n=406 pairs) with sample sizes >3.* ***C,*** *Population cluster analysis of P. falciparum microhaplotypes from dominant alleles in Zanzibar. Individual ancestry coefficients and optimum K value (K=3) are shown as inferred by rmaverick. Each vertical bar represents an individual haplotype and its membership to the three cluster are defined by the different colors. Black borders separate the five districts.*

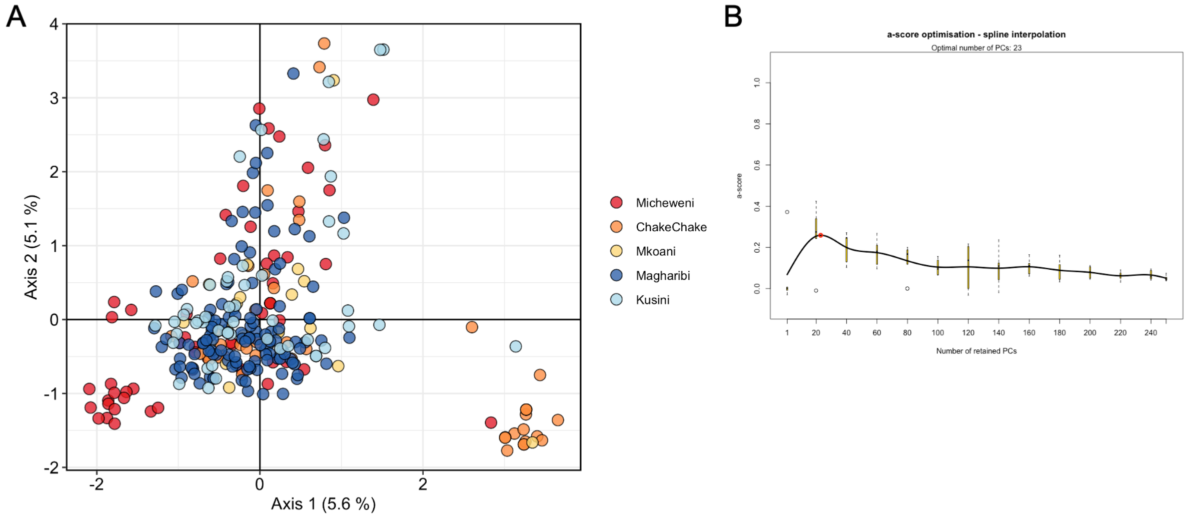


***Figure S6. A,*** *Principal components analysis (PCA) using allele frequency of the 28 microhaplotypes. The first two PCA components are shown (explaining 5.6% and 5.1% of variation in the dataset, respectively).* ***B,*** *Ascertaining the number of components to include in the discriminatory analysis of principal components (DAPC). Alpha-score optimization shows the optimal number of principal components (23 PCs) to retain for DAPC analysis based on the maximum a-score (red dot). These 23 PCs explain 49.8% of variation in the dataset.*

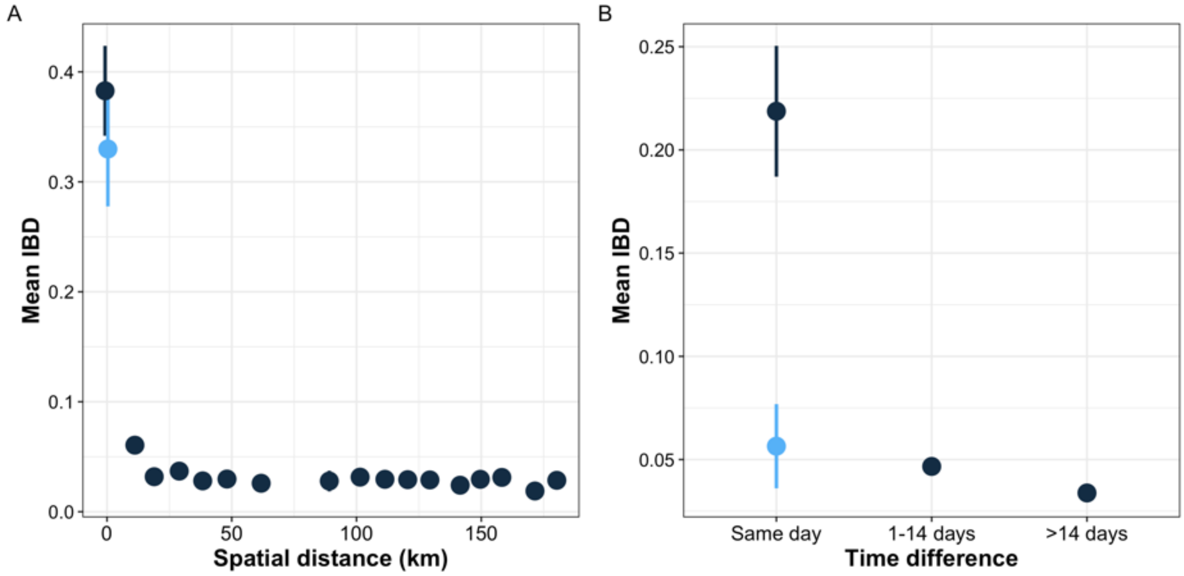


***Figure S7***. *Mean IBD by spatial distance and time difference.* ***A,*** *Spatial patterns in IBD. Mean IBD binned by the spatial distance in 10km intervals. Vertical lines indicate 95% confidence intervals. Blue dot indicates mean IBD where RACD pairs were excluded.* ***B,*** *Temporal patterns in IBD. Mean IBD of sample pairs collected on the same day, 1-14 days apart or >14 days apart. Vertical lines indicate 95% confidence intervals. Blue dot indicates mean IBD where RACD pairs were excluded.*

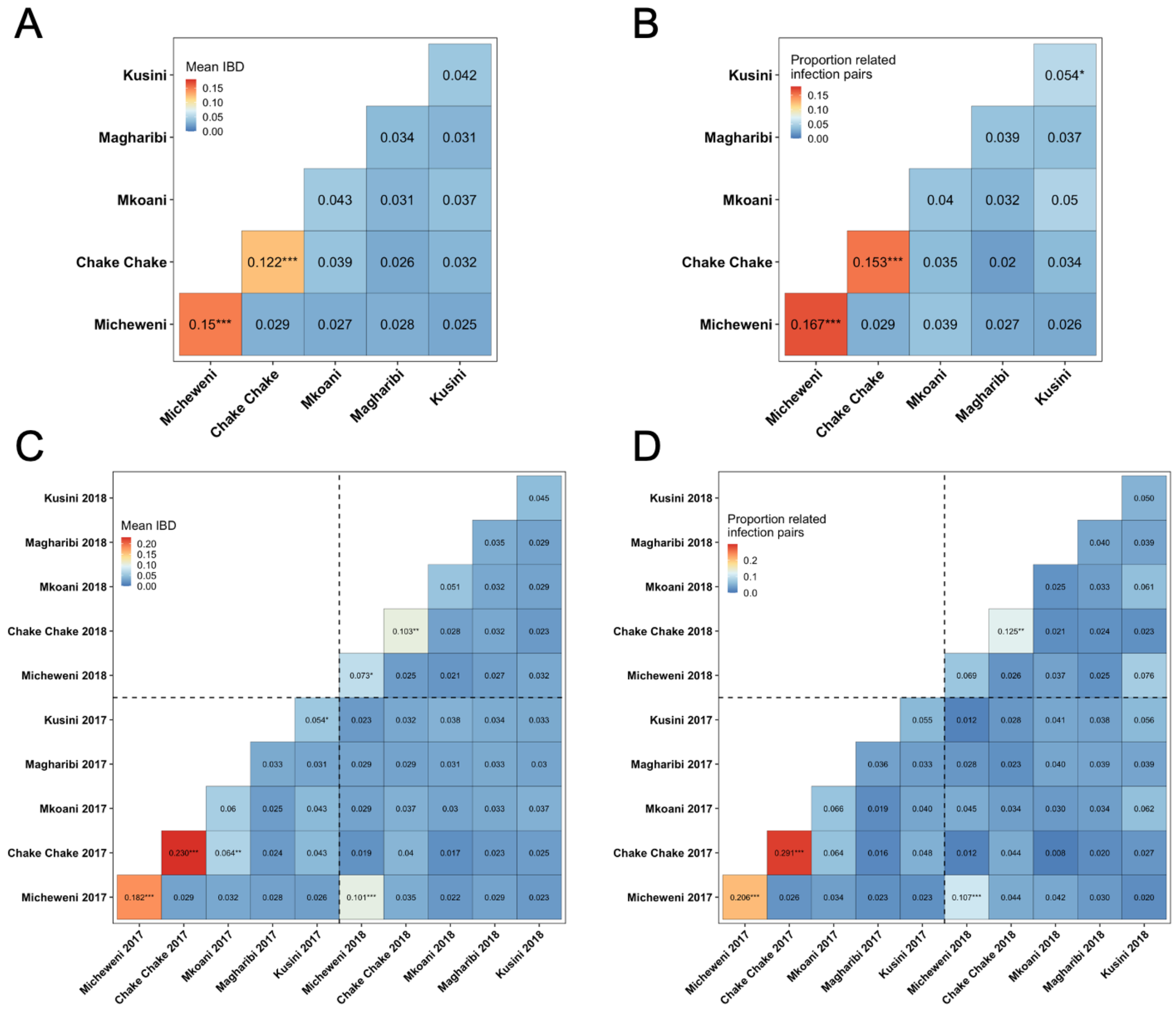


***Figure S8. A,*** *Mean IBD by districts.* ***B,*** *Proportion of related infection pairs by district.* ***C,*** *Mean IBD by districts stratified by year.* ***D,*** *Proportion of related infection pairs by district stratified by year. Asterisks correspond to the permutation test's p-value, P < 0.05 (*), P < 0.01 (**), and P < 0.001 (***).*

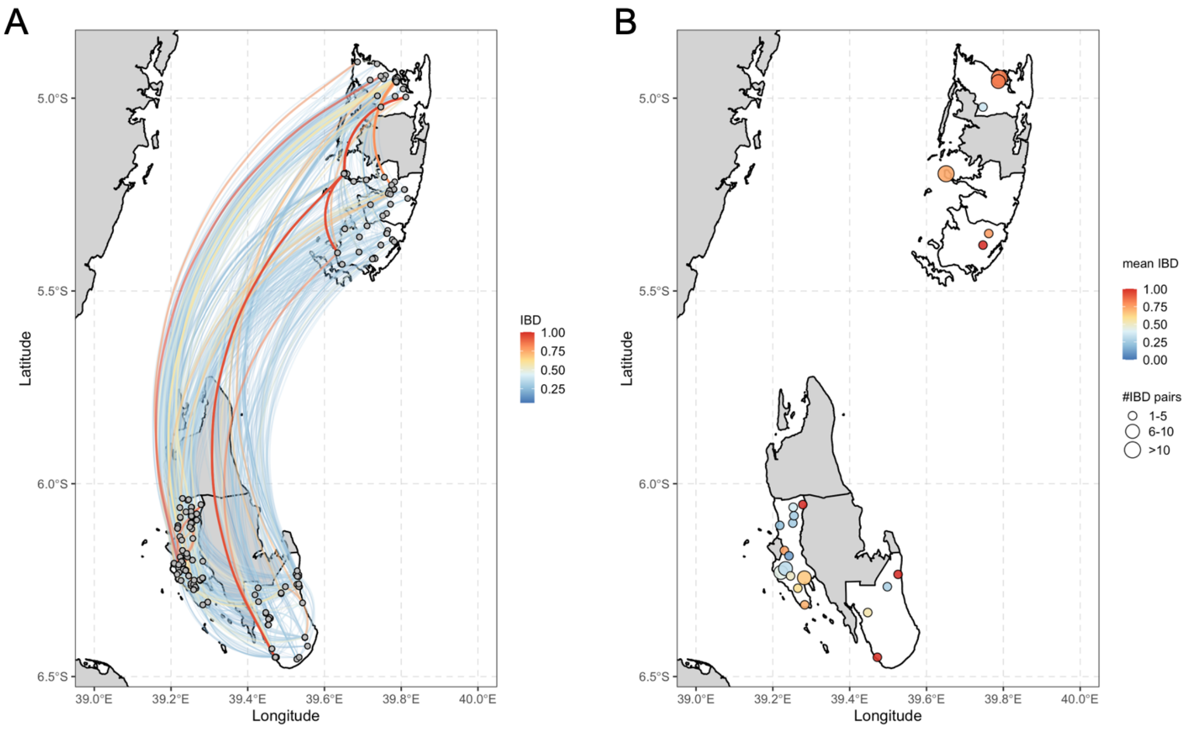


***Figure S9.*** *All significantly related pairs (n=1,703) (**A**) between shehia (n=1,335) and (**B**) within shehia (n=368). Within shehia mean IBD is higher on Pemba Island (0.78) than on Unguja Island (0.49).*

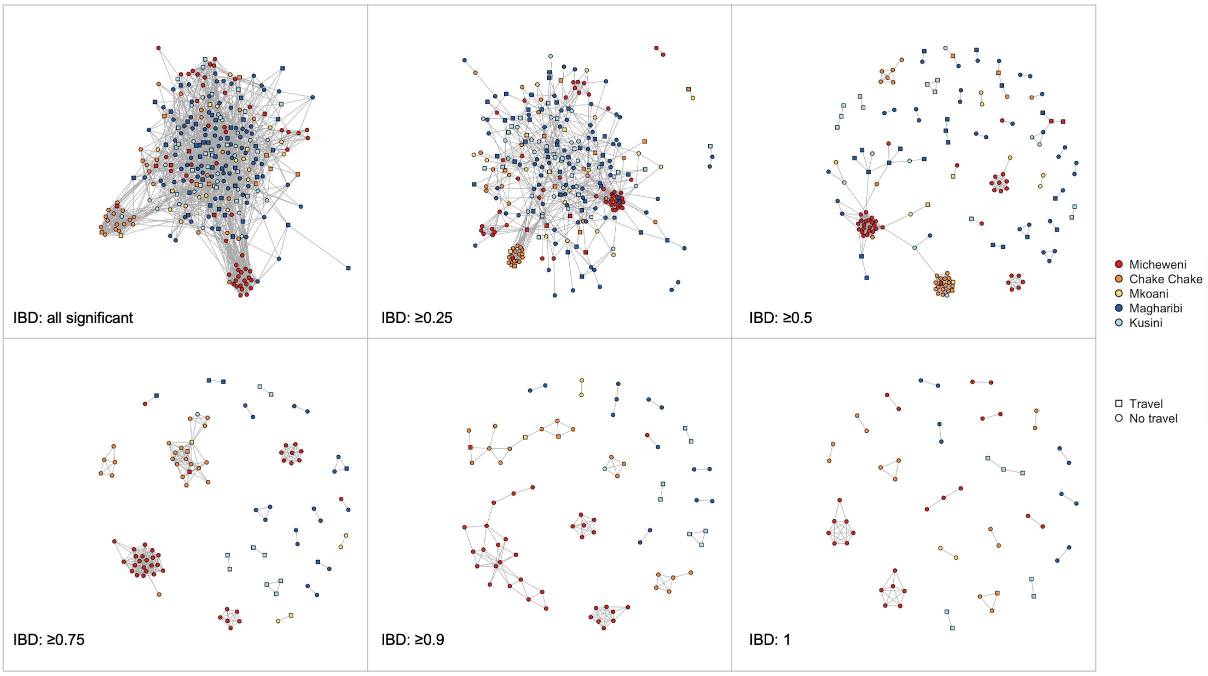


***Figure S10.*** *Network analysis of pairwise relatedness visualized using different thresholds of IBD and colored by district. Isolates that share less than the indicated IBD threshold with any other isolate are omitted from the network.*

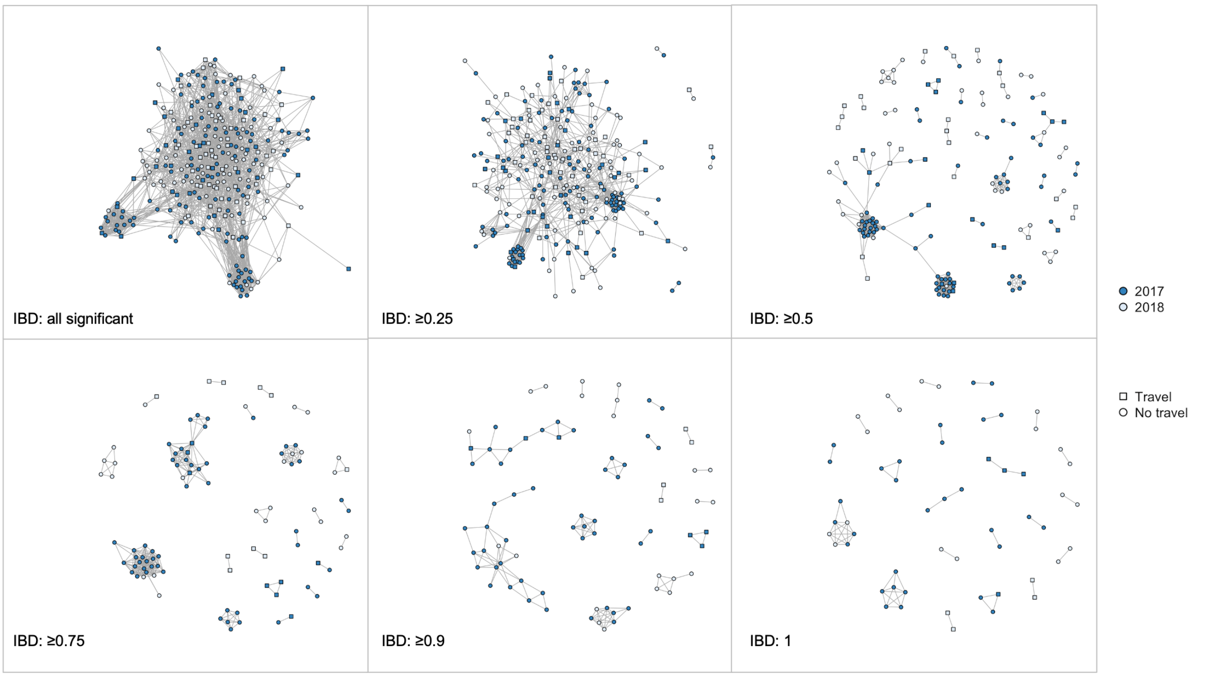


***Figure S11.*** *Network analysis of pairwise relatedness visualized using different thresholds of IBD and colored by year. Isolates that share less than the indicated IBD threshold with any other isolate are omitted from the network.*

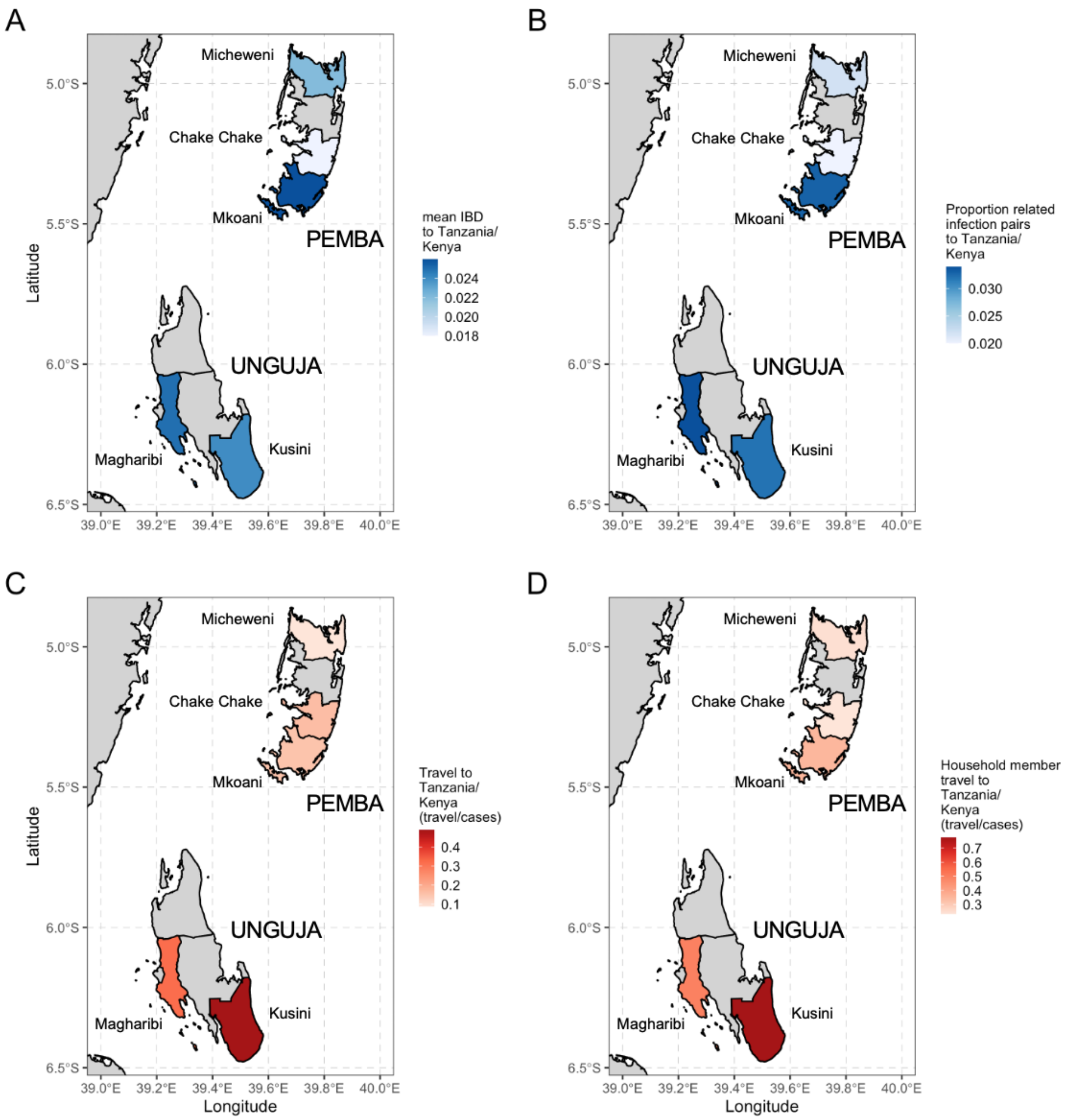


***Figure S12.*** *Map showing mean IBD and proportion of related infection of the five districts to mainland Tanzania and Kenya.* ***A****, Mean IBD of Micheweni (0.022), Chake Chake (0.018), Mkoani (0.026), Magharibi (0.025) and Kusini (0.024) from north to south.* ***B,*** *Proportion of related infection of Micheweni (0.022), Chake Chake (0.020), Mkoani (0.033), Magharibi (0.034) and Kusini (0.032).* ***C****, Proportion of people with genotyping data that reported travel to the mainland: Micheweni (8.8%), Chake Chake (16.3%), Mkoani (15.0%), Magharibi (32.2%) and Kusini (49.0%) from north to south.* ***D****, Proportion of household members that reported travel to the mainland from those individuals with genotyping data: of Micheweni (24.6%), Chake Chake (23.3%), Mkoani (35.0%), Magharibi (49.6%) and Kusini (77.6%) from north to south.*

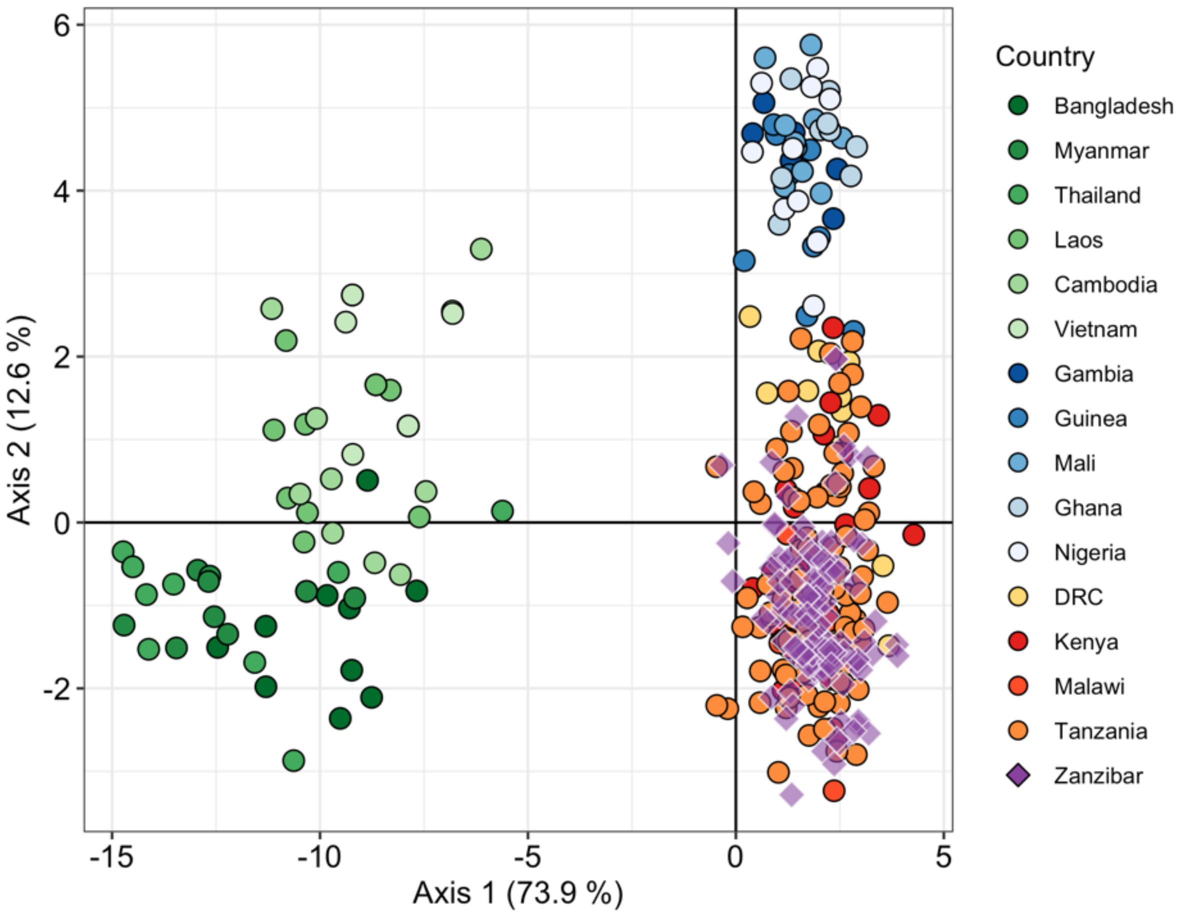


***Figure S13.*** *Genetic differentiation between global P. falciparum populations by discriminatory analysis of principal components (DAPC) using 28 microhaplotypes and 7 drug resistance loci. The first 48 components of a principal component analysis (PCA) are shown. Each point represents a single isolate (n = 374); colors indicate country of origin. Only monoclonal samples were used for this analysis.*

### Supplementary Tables

***Table S1.*** *Number of reads for whole blood and dried blood spot control samples.*

| Sample | Parasite density (per µL) | n | Median reads (IQR) [a] | Median reads per locus (IQR) [a] |
|---|---|---|---|---|
| **Whole blood** | | 20 | 491,561 (347,481 – 627,777) | |
| | 10,000 | 5 | | 6,830 (2,154 – 19,486) |
| | 1,000 | 5 | | 5,387 (1,351 – 19,354) |
| | 100 | 5 | | 5,237 (1,225 – 19,360) |
| | 10 | 5 | | 2,225 (860 – 8,659) |
| **Dried blood spots** | | 13 | 251,168 (116,148 – 460,167) | |
| | 1,000 | 3 | | 11,244 (5,229 – 27,684) |
| | 100 | 5 | | 2,408 (708 – 6,691) |
| | 10 | 5 | | 494 (133 – 1,769) |

[a] after trimming of low-quality reads.

***Table S2.*** *Detection of minority clone in mixed strain infections from cultured parasites at different ratios and parasite densities.*

| Strain mixtures | Abundance of minority clone | Parasite density (per µL) | Expected number of minority clones per µL to be detected | % of heterozygotes detected in minority clone (n/N) with cut-off [a] | | | | % of heterozygotes detected in minority clone (n/N) without cut-off [b] | | | |
|---|---|---|---|---|---|---|---|---|---|---|---|
| | | | | Rep 1 | Rep 2 | Rep 3 | Rep 4 | Rep 1 | Rep 2 | Rep 3 | Rep 4 |
| **3D7:FCB** | 40% | 1,000 | 400 | 100.0% (27/27) | 100.0% (27/27) | - | - | 100.0% (27/27) | 100.0% (27/27) | - | - |
| | 40% | 100 | 40 | 100.0% (27/27) | 100.0% (27/27) | - | - | 100.0% (27/27) | 100.0% (27/27) | - | - |
| | 40% | 10 | 4 | 63.0% (17/27) | 92.6% (25/27) | - | - | 63.0% (17/27) | 92.6% (25/27) | - | - |
| | 20% | 1,000 | 200 | 70.4% (19/27) | 70.4% (19/27) | - | - | 74.1% (20/27) | 77.8% (21/27) | - | - |
| | 20% | 100 | 20 | 66.7% (18/27) | 77.8% (21/27) | - | - | 77.8% (21/27) | 77.8% (21/27) | - | - |
| | 20% | 10 | 2 | 48.1% (13/27) | 66.7% (18/27) | - | - | 48.1% (13/27) | 70.4% (19/27) | - | - |
| **3D7:HB3** | 2% | 1,000 | 20 | 52.4% (11/21) | 61.9% (13/21) | 66.7% (14/21) | 71.4% (15/21) | 71.4% (15/21) | 71.4% (15/21) | 71.4% (15/21) | 71.4% (15/21) |
| | 1% | 1,000 | 10 | 28.6% (6/21) | 33.3% (7/21) | 33.3% (7/21) | 42.9% (9/21) | 47.6% (10/21) | 61.9% (13/21) | 66.7% (14/21) | 47.6% (10/21) |

[a] "With cut-off" meaning a within-host haplotype frequency of ≥1%.
[b] "No cut-off" meaning a within-host haplotype frequency of ≥0.1%.

***Table S3.*** *Number of samples included in further analyses (290/518) by districts. No significant difference in proportion of samples included by districts.*

| **Population** | **Samples included** | ***P value*** [a] | ***Chi-square statistic*** |
|---|---|---|---|
| **Districts** | | 0.6395 | 2.5285 |
| **Micheweni** | 59.3% (57/96) | | |
| **Chake Chake** | 50.0% (43/86) | | |
| **Mkoani** | 58.8% (20/34) | | |
| **Magharibi** | 57.9% (121/209) | | |
| **Kusini** | 52.7% (49/93) | | |

[a] *P*-value were two-tailed and computed using the chi-squared test for comparing the proportion of samples by district.

***Table S4.*** *Estimates of multilocus linkage disequilibrium (mLD) for P. falciparum populations in Zanzibar. $I_A^S$ = standardized index of association. The Monte Carlo method (100,000 permutations) was used to test the significance of mLD.*

| **Population** | **All infections** | | **Monoclonal infections** | |
|---|---|---|---|---|
| | **n** | **$I_A^S$ (*P* value)** | **n** | **$I_A^S$ (*P* value)** |
| **Pemba Island** | 51 | 0.0691 (<0.001) | 14 | 0.0847 (<0.001) |
| **Micheweni** | 25 | 0.1485 (<0.001) | 7 | 0.2682 (<0.001) |
| **Chake Chake** | 19 | 0.1402 (<0.001) | 6 | 0.1264 (<0.001) |
| **Mkoani** | 7 | 0.0056 (0.355) | 1 | NA |
| **Unguja Island** | 73 | 0.0077 (<0.001) | 14 | 0.0376 (<0.001) |
| **Magharibi** | 57 | 0.0054 (0.02) | 9 | -0.0022 (0.606) |
| **Kusini** | 16 | 0.0582 (<0.001) | 5 | 0.3437 (<0.001) |
| **All** | 124 | 0.0168 (<0.001) | 28 | 0.0356 (<0.001) |

***Table S5.*** *Mean IBD and proportion of related infection pairs in Zanzibar, stratified by island, district, shehia, RACD cluster, and year. P-values of each comparison were calculated by permutation test (100,000 permutations).*

| Comparisons | | Mean IBD | Fold change | *P value* | Proportion related infections | Fold change | *P value* |
|---|---|---|---|---|---|---|---|
| **Island** | | | 2.33 | < 0.0001 | | 2 | < 0.0001 |
| | **Pemba Island** | 0.07 | | | 0.08 | | |
| | **Unguja Island** | 0.03 | | | 0.04 | | |
| **District** | | | 2 | < 0.0001 | | 3 | < 0.0001 |
| | **Within** | 0.06 | | | 0.09 | | |
| | **Between** | 0.03 | | | 0.03 | | |
| ***Shehia*** | | | 2.25 | < 0.0001 | | 1.75 | < 0.0001 |
| | **Within** | 0.09 | | | 0.07 | | - |
| | **Between** | 0.04 | | | 0.04 | | - |
| **RACD cluster** | | | 3.25 | < 0.0001 | | 1.13 | < 0.0001 |
| | **Within** | 0.13 | | | 0.043 | | |
| | **Between** | 0.04 | | | 0.038 | | |
| **Year** | | | 1.33 | < 0.0001 | | 1.33 | < 0.0001 |
| | **Within** | 0.04 | | | 0.04 | | |
| | **Between** | 0.03 | | | 0.03 | | |
| | | | 1.66 | < 0.0001 | | 1.25 | 0.0003 |
| | **2017** | 0.05 | | | 0.05 | | |
| | **2018** | 0.03 | | | 0.04 | | |
| **Travel history** | | | 1.33 | 0.003 | | 1.66 | 0.0026 |
| | **Non-travelers** | 0.04 | | | 0.05 | | |
| | **Travelers** | 0.03 | | | 0.03 | | |

***Table S6***. *Mean IBD and proportion of related infection pairs between Zanzibar and mainland Tanzania/Kenya, stratified by island, district and travel/no travel. P values of each comparison between travel vs. non-travel were calculated by permutation test (100,000 permutations).*

| **Pairs** | **n pairs** | **Mean IBD** | ***P value*** | **Proportion related infections** | ***P value*** |
|---|---|---|---|---|---|
| **Zanzibar – Tanzania/Kenya** | 33,350 | 0.023 | - | 0.029 | - |
| **Zanzibar – Tanzania** | 23,200 | 0.024 | - | 0.031 | - |
| **Zanzibar – Kenya** | 10,150 | 0.022 | - | 0.024 | - |
| **Unguja Island – Tanzania/Kenya** | 19,550 | 0.025 | - | 0.034 | - |
| **Travel** | 7,245 | 0.025 | 0.5804 | 0.031 | 0.9366 |
| **No travel** | 12,305 | 0.025 | | 0.035 | |
| **Magharibi** | 13,915 | 0.025 | - | 0.034 | - |
| **Travel** | 4,485 | 0.026 | 0.2809 | 0.032 | 0.7717 |
| **No travel** | 9,430 | 0.025 | | 0.035 | |
| **Kusini** | 5,635 | 0.024 | - | 0.032 | - |
| **Travel** | 2,760 | 0.023 | 0.8187 | 0.028 | 0.9224 |
| **No travel** | 2,875 | 0.025 | | 0.037 | |
| **Pemba Island – Tanzania/Kenya** | 13,800 | 0.021 | - | 0.023 | - |
| **Travel** | 1,725 | 0.026 | **0.0041** | 0.026 | 0.3043 |
| **No travel** | 12,075 | 0.020 | | 0.023 | |
| **Micheweni** | 6,555 | 0.022 | - | 0.022 | - |
| **Travel** | 575 | 0.024 | 0.2227 | 0.017 | 0.8068 |
| **No travel** | 5,980 | 0.022 | | 0.023 | |
| **Chake Chake** | 4,945 | 0.018 | - | 0.020 | - |
| **Travel** | 805 | 0.024 | **0.0185** | 0.030 | 0.0561 |
| **No travel** | 4,140 | 0.017 | | 0.018 | |
| **Mkoani** | 2,300 | 0.026 | - | 0.033 | - |
| **Travel** | 345 | 0.037 | **0.0062** | 0.029 | 0.6917 |
| **No Travel** | 1,955 | 0.024 | | 0.034 | |

***Table S7.*** *Mean pairwise IBD and proportion of related infection pairs between samples from Zanzibar and several African countries.*

| Pairs with Zanzibar | n pairs | Mean IBD | Proportion related infections |
|---|---|---|---|
| **West Africa** | | | |
| **Gambia** | 2,900 | 0.017 | 0.024 |
| **Guinea** | 2,610 | 0.018 | 0.024 |
| **Mali** | 2,900 | 0.017 | 0.020 |
| **Ghana** | 2,900 | 0.016 | 0.019 |
| **Nigeria** | 2,900 | 0.016 | 0.019 |
| **Central Africa** | | | |
| **DRC** | 2,900 | 0.023 | 0.029 |
| **East Africa** | | | |
| **Kenya** | 10,150 | 0.022 | 0.025 |
| **Malawi** | 2,900 | 0.026 | 0.029 |
| **Tanzania** | 23,200 | 0.025 | 0.034 |
| **Zanzibar** | 41,905 | 0.040 | 0.045 |

***Table S8.*** *Frequency of mutations from drug resistance alleles in the population stratified by district.*

| **Gene** | **Mutation** | **Frequency% (n/N)** | | | | | ***P* value [a]** |
|---|---|---|---|---|---|---|---|
| | | **Micheweni** | **Chake Chake** | **Mkoani** | **Magharibi** | **Kusini** | |
| ***pfdhfr*** | C50R | 0% (0/38) | 0% (0/37) | 0% (0/18) | 0% (0/102) | 0% (0/36) | NA |
| | **N51I** | **100% (38/38)** | **97.3% (36/37)** | **83.3% (15/18)** | **96.1% (98/102)** | **97.2% (35/36)** | **0.048** |
| | C59R | 89.5% (34/38) | 89.2% (33/37) | 72.2% (13/18) | 86.3% (88/102) | 97.2% (35/36) | 0.1124 |
| | S108N | 100% (38/38) | 100% (37/37) | 100% (18/18) | 100% (102/102) | 100% (36/36) | NA |
| | S108T | 0% (0/38) | 0% (0/37) | 0% (0/18) | 0% (0/102) | 0% (0/36) | NA |
| | I164L | 0% (0/47) | 0% (0/38) | 0% (0/19) | 0% (0/110) | 0% (0/41) | NA |
| ***pfdhps*** | **K540E** | **95.3% (41/43)** | **47.2% (17/36)** | **68.8% (11/16)** | **92.6% (100/108)** | **83.3% (30/36)** | **0.0005** |
| | A581G | 0% (0/43) | 0% (0/36) | 0% (0/16) | 0% (0/108) | 0% (0/36) | NA |
| | A613T | 0% (0/43) | 0% (0/36) | 0% (0/16) | 0% (0/108) | 0% (0/36) | NA |
| | A613S | 0% (0/43) | 0% (0/36) | 0% (0/16) | 0% (0/108) | 0% (0/36) | NA |
| ***pfmdr1*** | N86Y | 2.4% (1/42) | 5.7% (2/35) | 0% (0/19) | 2.8% (3/106) | 7.3% (3/41) | 0.5967 |
| | Y184F | 68.5% (37/54) | 69.0% (29/42) | 63.2% (12/19) | 73.9% (85/118) | 48.9% (23/47) | 0.084 |
| ***pfmdr2*** | I492V | 37.3% (19/51) | 27.0% (10/37) | 44.4% (8/18) | 48.7% (56/115) | 47.6% (20/42) | 0.1759 |
| | T484I | 0% (0/51) | 0% (0/37) | 0% (0/18) | 0% (0/115) | 0% (0/42) | NA |
| ***pfk13*** | I543T | 0% (0/52) | 0% (0/40) | 0% (0/17) | 0% (0/114) | 0% (0/42) | NA |
| | R539T | 0% (0/52) | 0% (0/40) | 0% (0/17) | 0% (0/114) | 0% (0/42) | NA |
| | G538V | 0% (0/52) | 0% (0/40) | 0% (0/17) | 0% (0/114) | 0% (0/42) | NA |
| | N537I | 0% (0/52) | 0% (0/40) | 0% (0/17) | 0% (0/114) | 0% (0/42) | NA |
| | P527H | 0% (0/52) | 0% (0/40) | 0% (0/17) | 0% (0/114) | 0% (0/42) | NA |
| | Y493H | 0% (0/52) | 0% (0/40) | 0% (0/17) | 0% (0/114) | 0% (0/42) | NA |
| | A481V | 0% (0/52) | 0% (0/40) | 0% (0/17) | 0% (0/114) | 0% (0/42) | NA |

[a] All *P* values were 2-tailed and computed using the Pearson's Chi-squared test with simulated *P* value (based on 2000 replicates) for comparing the proportion of mutant alleles.
